# Two fundamental timescales organize human vocal behavior

**DOI:** 10.64898/2026.06.29.731229

**Authors:** Jing Wang, Honghua Chen, Yanchao Bi, Nai Ding

## Abstract

Humans produce a remarkable diversity of vocal behaviors throughout life, from innate sounds such as cry, laughter, and babble to learned, sophisticated sequences such as speech and song. Whether these diverse vocalizations share basic principles in their temporal organization remains unknown. Here, we show that, despite their distinct functions and extraordinary cultural variation, these vocalizations consistently occupy two characteristic rhythmic regimes supporting distinct functionalities. Corpus analyses revealed that, by approximately three years of age, the acoustic rhythms of speech and song settle into two distinct frequency ranges: one between 4 and 8 Hz, the other below 2 Hz. Critically, these two temporal regimes are already evident in infant cry, laughter, and babble, prior to the emergence of language and song. Sensorimotor synchronization experiments, where participants vocalized or tapped to sound sequences presented at different rates, revealed that individual vocal production exhibited maximal resonance between 4 and 8 Hz, whereas synchronization across individuals was strongest below 2 Hz. Together, these findings identify two functionally distinct sensorimotor timescales that organize diverse human vocal behavior: a faster timescale supporting stable individual vocal production and a slower timescale facilitating interpersonal synchronization.

## Introduction

Humans produce diverse vocalizations throughout life, ranging from innate sounds such as cry and laughter to culturally learned complex sequences such as speech and song (Fig. 1). Because these vocalizations are produced by the same articulatory system and perceived by the same auditory system, they may share intrinsic constraints imposed by the sensorimotor system. An influential hypothesis holds that the human sensorimotor circuitry for speech production and perception operates at a characteristic timescale between centered around 5 Hz (Poeppel & Assaneo, 2020). This hypothesis, referred to as the single-timescale hypothesis, explains the common speech rhythm observed across languages: the acoustic rhythm of speech, characterized by the modulation spectrum, peaks between 4 and 8 Hz (Ding et al., 2017). This rhythm has also been linked to theta-band (4–8 Hz) neural oscillations in sensory and motor cortices (Ghitza, 2012; Giraud & Poeppel, 2012; Poeppel & Assaneo, 2020) and may have evolutionary precursors in nonhuman primates (Bergman et al., 2019; Ghazanfar et al., 2012).

**Fig. 1.**
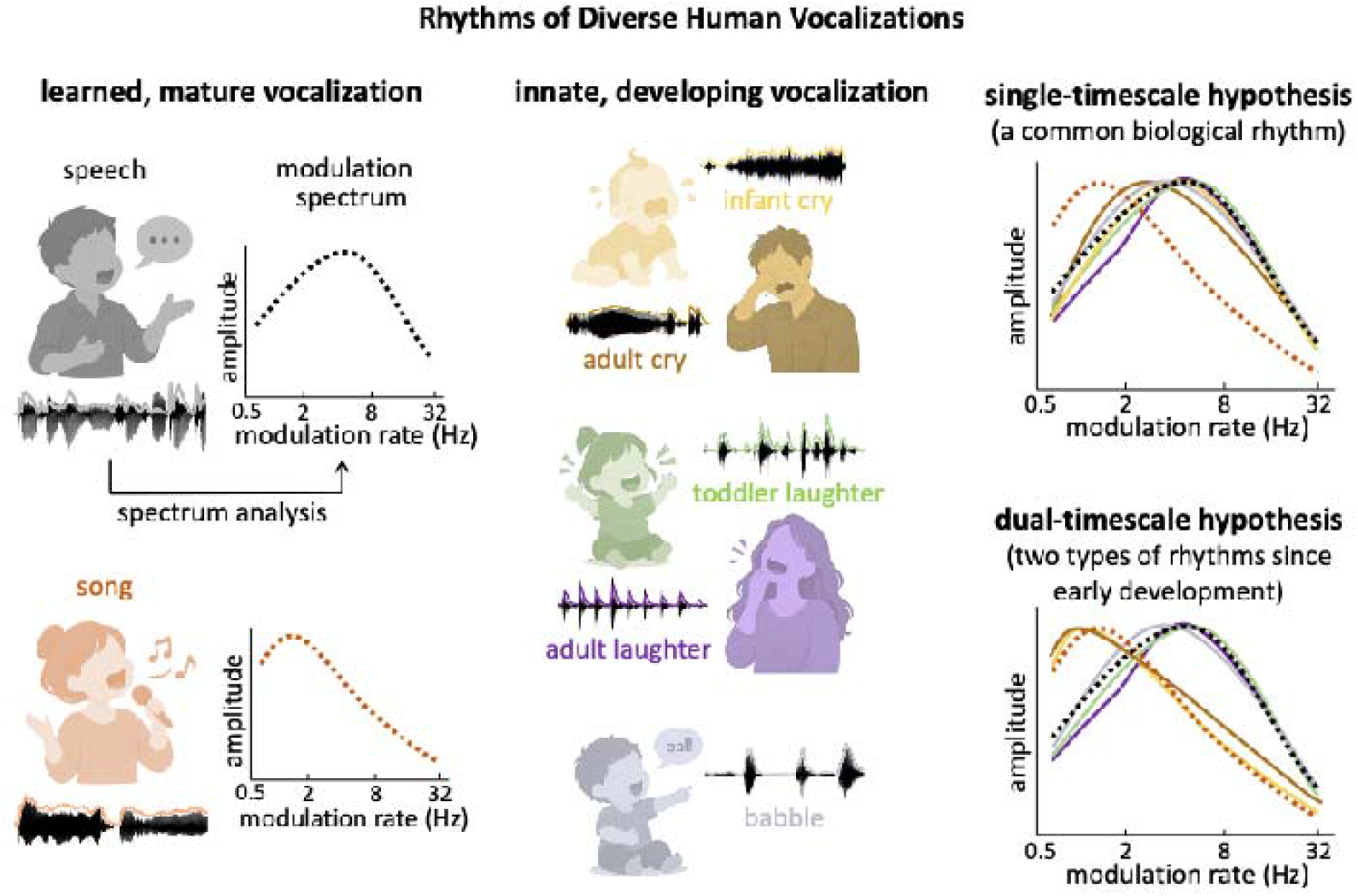
Rhythms of diverse human vocalizations. The acoustic rhythm of a sound can be characterized by the modulation spectrum. An influential hypothesis posits that the dominant 4-8 Hz speech rhythm reflects the characteristic frequency of the human vocal production/perception system (Poeppel & Assaneo, 2020). According to this single-timescale hypothesis, it is possible that most innate vocalizations, which share the same biological constraints, share the same rhythm. The rhythm of song, however, differs from the rhythm of speech. If the song rhythm represents another characteristic frequency of the human vocal system, innate vocalizations may adopt either characteristic frequency, i.e., the dual-timescale hypothesis. Representative vocalizations are shown with their waveform (black), temporal envelope (colored lines), and schematic modulation spectra.

A potential challenge to the single-timescale hypothesis comes from music. Music, including song, is universally observed across cultures and exhibits a slower acoustic rhythm. When characterized by the modulation spectrum, the acoustic rhythm of music peaks below 1 Hz, rather than between 2 and 8 Hz as in speech (Albouy et al., 2024; Ding et al., 2017). One likely cause for the rhythmic difference between speech and music is their distinct social functions. Speech primarily serves to convey information (Fedorenko et al., 2024), whereas music fosters social bonding and coordinates joint movement (Mehr et al., 2018, 2019; Savage et al., 2021; Singh & Mehr, 2023; Yurdum et al., 2023). Consistent with this functional division, music is often performed by a group in synchrony, while speech is typically delivered by an individual (Shilton et al., 2023; Stivers et al., 2009). Previous studies have shown that humans can better synchronize their movements with rhythms below 2 Hz (Repp & Su, 2013).

According to the single-timescale hypothesis, music production operates outside the characteristic timescale of the sensorimotor system and might therefore require additional effort. Another possibility, however, is that the sensorimotor system contains two characteristic timescales, i.e., the dual-timescale hypothesis, allowing speech and song to exploit different sweet spots of the system and better serve their social functions. The single- and dual-timescale hypotheses lead to distinct predictions (Fig. 1 and Fig. 2a). First, the intrinsic timescale(s) of the sensorimotor system should also constrain innate vocalizations. If 2–8 Hz is the only intrinsic timescale, it should be pervasively observed in innate vocalizations; if both 2–8 Hz and <1 Hz timescales exist, the acoustic rhythms of innate vocalizations should bifurcate. Second, early vocalizations are likely to be constrained by the intrinsic timescale(s) of the sensorimotor system. If song rhythm originates from speech rhythm and diverges from it under social influences, it should change gradually over development. Third, the intrinsic timescale(s) should also be observable in basic sensorimotor tasks that do not involve stereotyped vocal productions. Furthermore, given the social functions of speech and music, if two timescales exist, they may be separately optimal for individual production and interpersonal synchronization.

**Fig. 2.**
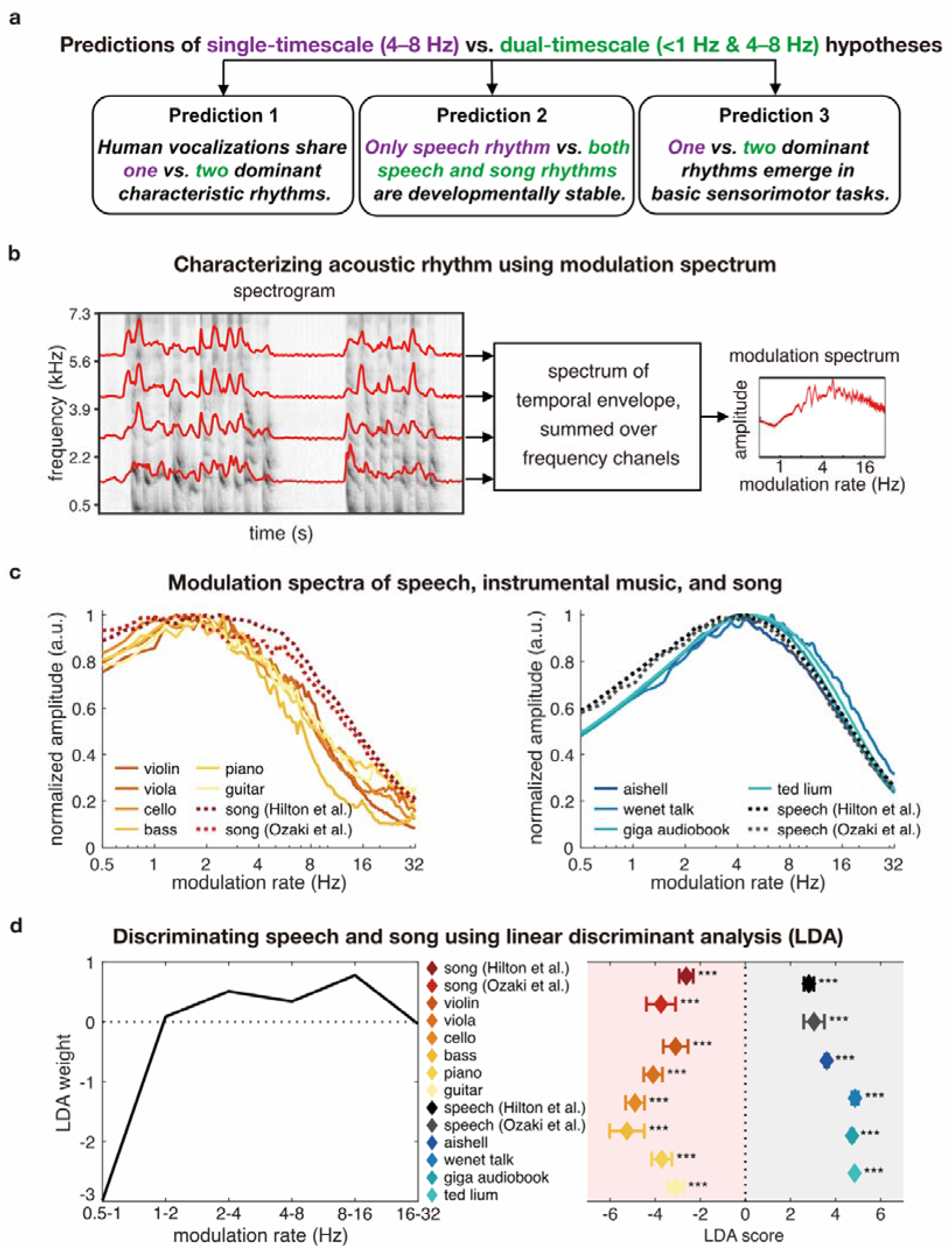
Predictions and analysis methods. (a) Predictions of the single-timescale vs. dual-timescale hypotheses. (b) Schematic illustration of modulation spectrum analysis, which extracts the temporal envelope from each cochlear frequency channel and applies a spectral analysis of the temporal envelope. (c) Modulation spectra of music (left) and speech (right). Each line represents one corpus. Solid lines show speech and instrumental music corpora analyzed in Ding et al. (2017). Dotted lines show speech and song recordings from two cross-lingual corpora (Hilton et al., 2022; Ozaki et al., 2024). (d) Linear discriminant analysis (LDA) was trained to discriminate speech and song using the two cross-lingual corpora and applied to the speech and instrumental music corpora used in Ding et al. (2017). The left panel shows the LDA weights at each modulation frequency. The right panel shows LDA scores for each corpus. Diamonds indicate group means, and horizontal bars indicate ±1 SEM across speakers/songs. Asterisks denote significance levels of LDA scores relative to zero (*p < 0.05, **p < 0.01, ***p < 0.001; two-sided t tests with FDR correction).

To test the first two predictions, we used the modulation spectrum framework (Fig. 2b) to analyze the acoustic rhythms of speech, song, and innate vocalizations (cry, laughter, babble), and to track how these rhythms change over development. To test the third prediction, we designed sensorimotor experiments in which participants tracked rhythmic or arrhythmic sensory sequences presented at different rates. We additionally quantified inter-participant synchronization to examine whether the rhythms that dominate individual production differ from those that maximize interpersonal alignment.

## Results

### Speech and song modulation spectra as baselines

We first characterized the modulation spectra (see Fig. 2b) of speech and the unaccompanied song, and compared them with the modulation spectra of speech and instrumental music reported in a previous study (Ding et al., 2017). Although the unaccompanied song was produced vocally while instrumental music was produced through manual actions, their modulation spectra showed a similar shape (Fig. 2c). To quantify the difference between speech and song and to provide a baseline for studying innate vocalizations, we projected the modulation spectrum into a one-dimensional linear discriminant analysis (LDA) score that was trained to classify speech and song. The LDA was an interpretable linear method and the weights it learned contrasted the power below 1 Hz and the power above 2 Hz. The LDA trained to classify speech and song also generalized to independent corpora of speech and instrumental music. Across corpora, speech samples yielded positive LDA scores and music (including song and instrumental music) yielded negative LDA scores (Fig. 2d). These LDA scores significantly differed from zero (see Table S1), and their polarity provided a generalizable measure of whether a modulation spectrum was more speech-like or song-like.

### Dual timescales in innate vocalizations

We next analyzed the modulation spectra of innate vocalizations, i.e., cry, babble, and laughter, and compared them with the modulation spectra of speech and song (Fig. 3a–c). Based on multilingual corpora, we found that cry showed song-like modulation spectra, whereas babble and laughter showed speech-like modulation spectra. To quantify the effects, we analyzed innate vocalization corpora using the LDA classifier trained to distinguish adult speech and song (Fig. 3d–f). The LDA scores of cries were generally below 0, confirming alignment with adult song. In contrast, the LDA scores of babbles and laughter were above 0, confirming alignment with adult speech (Fig. 3d–f; see Table S3 for statistics). These findings suggest innate vocalizations show both speech-like rhythms and music-like rhythms.

**Fig. 3.**
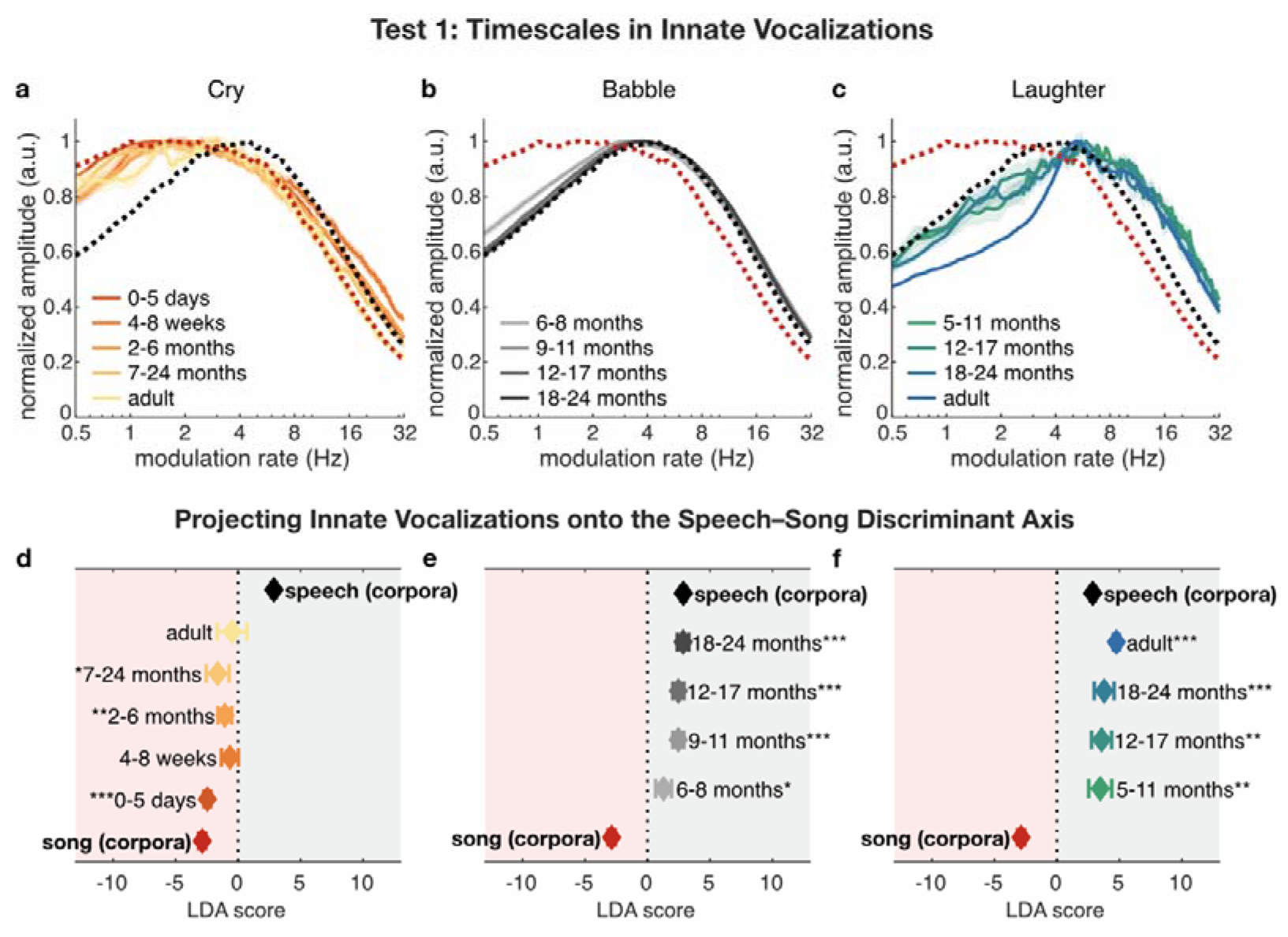
Two characteristic rhythms in cry, babble, and laughter. (a-c) Modulation spectra of cry (left), babble (middle), and laughter (right). Reference modulation spectra of speech and song (Hilton et al., 2022; Ozaki et al., 2024) are shown as dotted lines. Shaded regions indicate ±1 SEM across participants. (d-f) LDA scores for the modulation spectrum of cry (left), babble (middle), and laughter (right). Diamonds indicate group means, and horizontal bars indicate ±1 SEM across participants. Asterisks denote significance levels of LDA scores relative to zero (*p < 0.05, **p < 0.01, ***p < 0.001; two-sided t tests with FDR correction).

### Speech and song rhythms diverge early in development

Analyses of innate vocalizations revealed two characteristic rhythms, which are evident even before the emergence of speech and song. Next, we examined whether the acoustic rhythms of speech and song emerge early in development and align with these pre-existing rhythms. We analyzed the modulation spectra of speech and song recorded from speakers between 2 and 18 years of age (Fig. 4a–c). The modulation spectra were generally preserved across ages: song exhibited stronger low-frequency modulations below 2 Hz, whereas speech exhibited stronger modulations between 4 and 8 Hz. The peak frequencies of speech and song significantly diverged by age 3 (Fig. S1; see Table S2 for statistics).

**Fig. 4.**
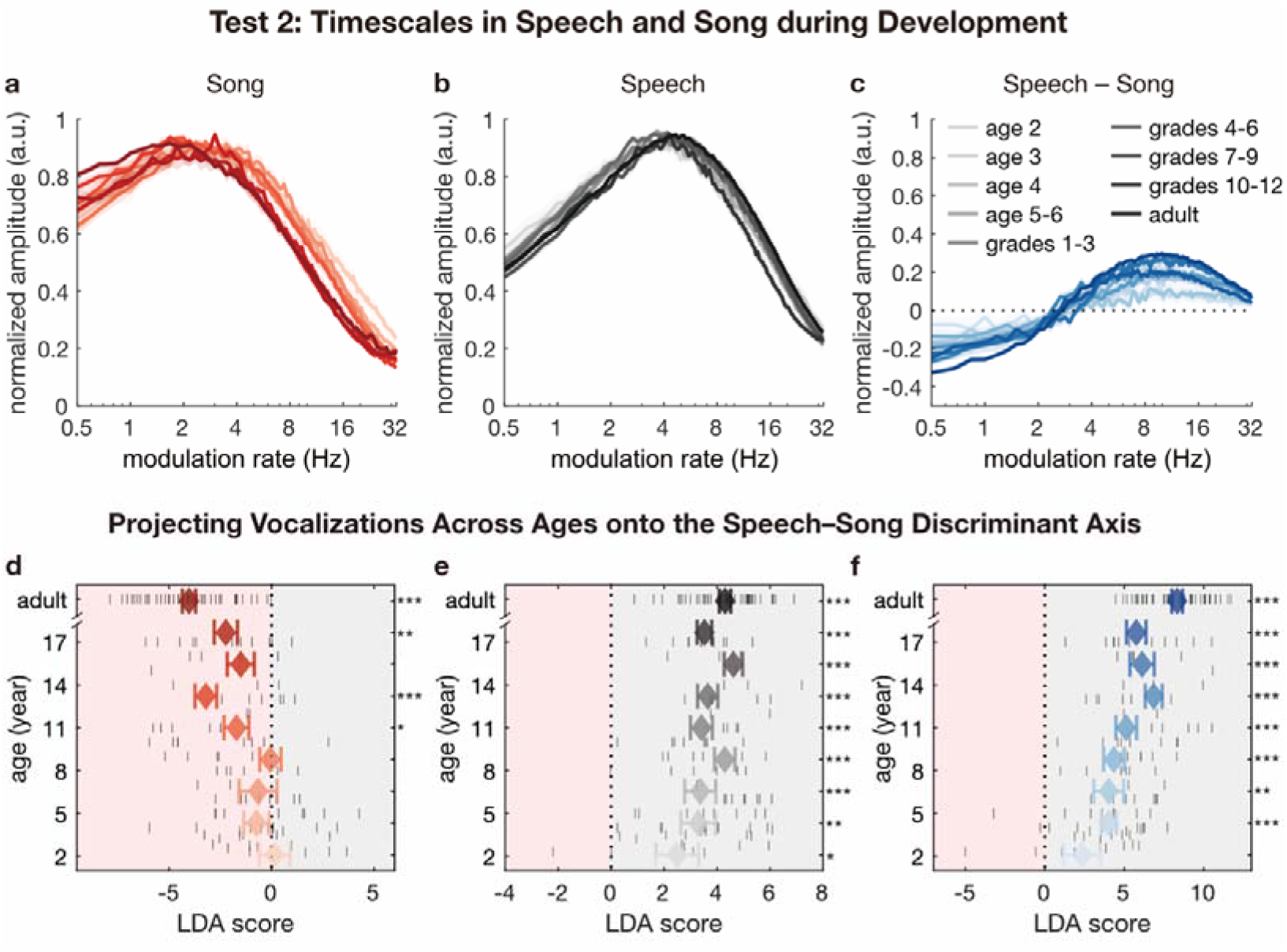
Early divergence between speech and song rhythms. (a-c) Group-averaged modulation spectra of song (left), speech (middle), and their difference (speech minus song; right) across age groups. Colors denote different age groups. Shaded regions indicate ±1 SEM across participants. (d-f) LDA scores for modulation spectra of song (left), speech (middle), and their difference (right). Diamonds indicate group means, horizontal bars indicate ±1 SEM across participants, and short lines indicate individual data points. The dotted vertical line indicates an LDA score of zero. Asterisks denote significance levels of LDA scores relative to zero (*p < 0.05, **p < 0.01, ***p < 0.001; two-sided t tests with FDR correction).

To further confirm that the two characteristic rhythms were already reliably expressed in early speech and song, we tested whether the LDA classifier trained to distinguish adult speech and song could distinguish speech and song produced at younger ages (Fig. 4d–f). Across all ages, the LDA scores for speech were consistently above 0 (Fig. 4e), and the LDA scores for song were generally below 0 and became significant in children older than 6 years (Fig. 4d). The difference between speech and song LDA scores was already significantly above 0 at age 3 (Fig. 4f). Statistics are provided in Table S3. These findings reveal that speech and song rapidly converge on the faster and slower timescales, respectively.

### Dual timescales in stimulus-synchronized sound production

The previous results suggested two characteristic timescales in human vocalizations. Next, we asked whether these two timescales can spontaneously emerge during sensorimotor synchronization. In the experiments, participants were asked to produce a sound sequence in synchrony with an external rhythm spanning rates faster or slower than the typical rates in speech and music. In particular, we separately analyzed the timescale preferred by an individual participant using the modulation spectrum and the timescale that may better synchronize a group of participants using the inter-participant phase coherence.

In one condition, the external rhythm was a periodic sequence of noise bursts. It formed a steady beat in each trial and the beat rate was between 1 and 7 Hz (Fig. 5a). Participants were asked to produce a syllable in synchrony to the external beat, and we then computed the modulation spectrum of their vocalization (Fig. 5c). At each external beat rate, the modulation spectrum showed a clear spectral line at the beat rate and its harmonics, confirming that the participants produced a periodic sequence in sync with the external beat. The response amplitude, however, was rate dependent. The envelope of the modulation spectra at different beat rates peaked at 6.01 Hz (Fig. 5e).

**Fig. 5.**
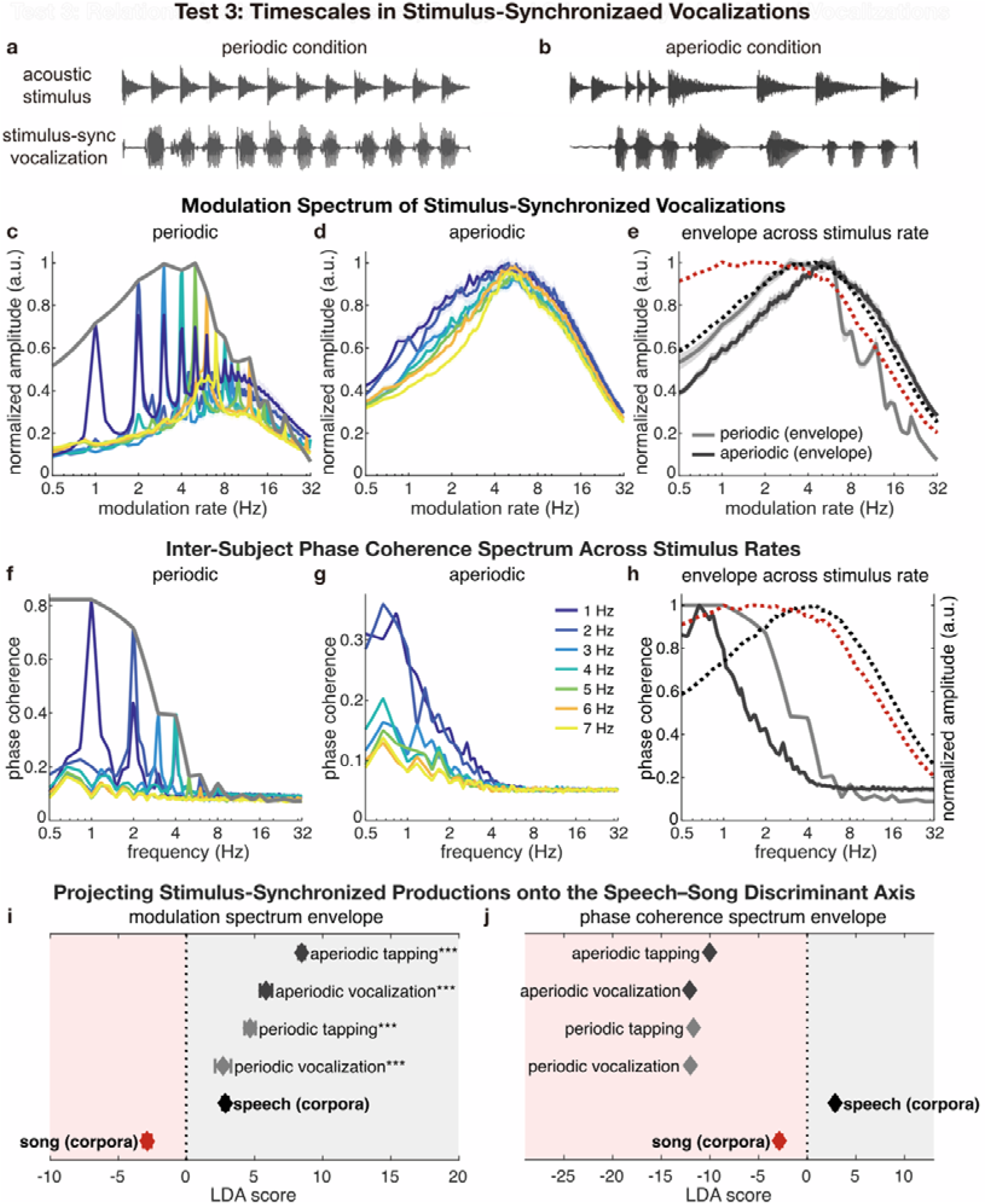
Two characteristic rhythms emerge during sensorimotor tracking. (a-b) Waveform of example stimulus and stimulus-synchronized vocalization from a representative participant under the periodic (a) and aperiodic (b) stimulus conditions. (c-d) Modulation spectrum of stimulus-synchronized vocalization at different stimulus rates (1–7 Hz) under the periodic (c) and aperiodic (d) conditions. Colors denote different stimulus rates. Shaded regions indicate ±1 SEM across participants. (e) Envelope of modulation spectra across different rates in the periodic (light gray) and aperiodic (dark gray) conditions. Shaded regions indicate ±1 SEM across participants. Reference modulation spectra of speech and song (Hilton et al., 2022; Ozaki et al., 2024) are shown as dotted lines. (f-g) Inter-participant phase coherence spectra of stimulus-synchronized vocalization. (h) Envelope of phase coherence spectra across different rates. (i-j) LDA scores for the envelope of modulation spectrum (i) and phase coherence spectrum (j). Diamonds indicate the mean, and horizontal bars indicate ±1 SEM across participants. Asterisks denote significance levels of LDA scores relative to zero (*p < 0.05, **p < 0.01, ***p < 0.001; two-sided t tests with FDR correction).

In the other condition, the external rhythm was aperiodic and the mean rate of noise bursts varied between 1 and 7 Hz (Fig. 5b). The modulation spectrum of the vocalization produced by the participants generally peaked between 4 and 8 Hz (Fig. 5d), and the envelope spectrum peaked at 4.98 Hz (Fig. 5e). These results indicated that, even when the participant attempted to produce a rhythm at various rates, their vocalization still reliably showed a resonance between 4 and 8 Hz, consistent with the acoustic rhythm of speech. Furthermore, to test whether this effect was specific to the effector, we also asked the participants to tap to the external rhythm and found very similar results (Fig. S2).

Next, we analyzed the synchrony among participants using the inter-participant phase coherence at each modulation frequency (Fig. 5f–g; Fig. S2). High phase coherence indicated that the participants produced sounds in synchrony, which could more effectively sum up if participants were to produce the sounds together. The envelope of the phase coherence spectrum peaked below 2 Hz, for both stimulus-synchronized vocalization and tapping (Fig. 5h; Fig. S2). The LDA analysis also confirmed that the LDA scores of the modulation spectrum envelope were significantly above 0 (Fig. 5i). The LDA scores for phase coherence spectrum envelopes were consistently below 0 (Fig. 5j). Exact statistics are provided in Table S3. These results all demonstrate two distinct sensorimotor timescales: a faster timescale between 4 and 8 Hz that dominates individual vocal production, and a slower timescale below 2 Hz that maximizes interpersonal synchronization.

## Discussion

The current study demonstrates that diverse human vocal behaviors are organized based two characteristic timescales, i.e., a faster timescale between 4 and 8 Hz and a slower timescale below 1 Hz. The timescales appear to be intrinsic properties of the vocal production system: they are present in innate vocalizations such as cry and laughter, appear in speech and song rhythms early in life, and emerge spontaneously when participants attempt to synchronize to external rhythms at different rates. Functionally, the sensorimotor synchronization experiment reveals that the faster timescale aligns with the resonance property of the individual sound production system, while the slower timescale aligns with the frequency range in which participants can better synchronize with each other.

### Acoustic rhythms of innate vocalizations

Rhythm is a fundamental feature of sound and is traditionally studied within separate domains such as speech and music (Patel, 2008). It has been proposed that an integrative and comparative approach that jointly considers the rhythms of speech, music, and animal vocalizations may shed new light on the biological and evolutionary bases of rhythm production and perception (Kotz et al., 2018; Ravignani, 2014). Here, we analyzed the acoustic rhythms of speech, music, and innate vocalizations, including cry, babble, and laughter, under the same framework of modulation spectrum. We observed that cry show a modulation spectrum that is similar to the modulation spectrum of music (Fig. 3a), consistent with a recent study showing that the modulation spectrum of newborn cry peaks around 2 Hz (Ruan et al., 2025). The modulation spectrum of laughter, in contrast, peaks around 6 Hz, more similar to speech (Fig. 3b), consistent with previous reports that human laughter consists of rapid acoustic bursts with rates comparable to speech syllables (Mowrer et al., 1987). The modulation spectrum of babble is also consistent with that of speech (Fig. 3c), consistent with previous findings that babbling exhibits syllable-like structure and develops continuously into speech (Locke, 1989). The current study integrates previous findings by consistently characterizing all sound categories using modulation spectrum analysis. The results suggest that the acoustic rhythms of speech and music can already be produced by infants (during crying and laughter) before they can produce connected (and therefore rhythmic) speech or music. In other words, the infant brain may be “rhythmically ready” within the first year of life.

Note that although cry and laughter are universal across human cultures, cry is influenced by the language prosody starting from early language exposure (Abboub et al., 2016; Mampe et al., 2009; Martinez-Alvarez et al., 2023; Nazzi et al., 1998). The language-specific prosodic patterns, however, are not strongly captured by the modulation spectrum. Indeed, different spoken languages, including languages falling into different prosodic categories such as English and French, show highly similar modulation spectra (Ding et al., 2017). In other words, the perceptual prosodic patterns of speech are not primarily rooted in the low-level acoustic features but instead may rely on phonetic features (Ramus et al., 2000). The current study, however, focuses on the acoustic rhythms of sounds, while the internal dynamical properties of the brain can further constrain how rhythms are perceived (Barbero et al., 2025; Jacoby et al., 2024; Jacoby & McDermott, 2017).

### Two timescales in vocal production

In a dynamical system, characteristic timescales can manifest in both spontaneous activity and in the response to an external input (Oppenheim et al., 1997). A simple dynamical system such as a spring has a single characteristic frequency. Both spontaneous oscillations and forced oscillations are strongest at the characteristic frequency. More complex systems may operate across multiple characteristic timescales to support distinct functional behaviors (Buzsáki & Llinás, 2017; Kelso, 1995; Poeppel, 2003). In the human brain, there may be partly separated neural pathways for speech and music processing (Norman-Haignere et al., 2015, 2022) and even the rhythm system per se engages multiple cortical and subcortical areas (Kotz et al., 2018), which may be wired in different ways to generate different characteristic frequencies. Here, we treat innate vocalizations (cry and laughter) as spontaneous activity of the human sound production system, and the sensorimotor synchronization experiment characterizes how the system responds to an external force. The two lines of evidence converge, both suggesting that the human sensorimotor system involved in vocal production possesses two intrinsic timescales.

On top of the sound categories we analyzed, the spontaneous tapping rate is also well known to be near 2 Hz (Repp, 2005; Repp & Su, 2013), although it is strongly influenced by musical experience (Drake et al., 2000). Integrating the results from different sound categories, it is clear that the 4-8 Hz and below 2 Hz rhythms are observed in sounds produced by different effectors. The 4-8 Hz rhythm is observed in oral speech and stimulus-synchronized vocalization (Fig. 5e), as well as in stimulus-synchronized tapping (Fig. S2). The below-2-Hz rhythm is observed in both orally produced song and cry (Fig. 3c), as well as in instrumental music (Fig. 2c) and spontaneous finger tapping (Fig. S2). These results suggest that the rhythms are generated in the central sensorimotor system instead of reflecting effector-specific timescales.

### Functional division between the two timescales

The sensorimotor experiment clearly demonstrates that individual participants most strongly produce a rhythm between 4 and 8 Hz. In the same experiment, however, the rhythms produced by the different participants are better synchronized below 2 Hz. Therefore, although individual participants produce a strong rhythm between 4 and 8 Hz, this fast rhythm is not aligned across participants and therefore cannot effectively sum up in a crowd. In contrast, the slow rhythm below 2 Hz can more effectively synchronize across individuals within a group. The slower sound rhythm can better serve the purpose of synchronizing movements, including collective behaviors such as dancing (Scheurich et al., 2018). More broadly, this synchronization may extend to the synchronization of emotion and attention (Fritz et al., 2009; Keller et al., 2014). A similar dissociation between the characteristic frequency of an individual and the characteristic frequency of a group have also been observed in other domains (Maurer & McNaughton, 2007; Sloin et al., 2024), and usually slower temporal dynamics are observed in larger networks (Hancock et al., 2025). Among human vocalizations, song and cry are the categories that can most effectively synchronize attention and emotional systems. Song is also the category that most likely engage synchronous production from multiple individuals (Mehr et al., 2019; Savage et al., 2015). These social needs may explain why song and cry adopt the slower rhythm. In contrast, speech is usually produced by an individual, and laughter is a social signal that is not always intended to be heard (Winkler & Bryant, 2021), which may explain why they adopt the faster rhythm to efficient production.

### Summary

In summary, our findings support the existence of two functionally distinct timescales that organize human vocal behavior. The presence of the two timescales in pre-linguistic vocalizations and in sensorimotor synchronization experiments suggest that they reflect fundamental properties of the human sensorimotor system. Speech and music may differentially recruit these two rhythms to serve the purposes of communication and interpersonal synchronization, respectively.

## Materials and Methods

### Audio Samples and Stimulus

**Speech and Music (adults):** We analyzed two cross-lingual corpora consisting of speech and song recordings from adults (Hilton et al., 2022; Ozaki et al., 2024) as well as the speech and instrumental music corpora used in a previous study (Ding et al., 2017).

**Cry, Babble, and Laughter:** We analyzed cry, babble, and adult laughter based on publicly available corpora (Davis & MacNeilage, 1995; Gong et al., 2022; Kern et al., 2009; Lintfert, 2010; Lockhart-Bouron et al., 2023) (Table S4). Toddler laughter was recorded (N = 27, aged 5 to 24 months; Table S5) through recruitment advertisements distributed via social media platforms. Recordings were collected by caregivers at home. All recordings were screened by the first author, and non-laughter portions were replaced with silence of the same duration. The experimental procedures were approved by the Research Ethics Committee of the College of Biomedical Engineering and Instrument Sciences, Zhejiang University (No. 2025–5). Parents received an information sheet describing the study procedures and could decline participation by returning an opt-out form. Participants received monetary compensation for participation.

### Recordings

**Speech and Song Recording (children):** We collected spontaneous speech and song recordings from children through recruitment advertisements distributed via social media platforms. Recordings were collected by caregivers. We received both speech and song recordings from 100 children (61 females, aged 2 to 17 years; Table S6). We received only speech or song recordings from 14 additional children, who were therefore excluded from the analysis. The content of speech and song was not restricted. The experimental procedures were approved by the Research Ethics Committee of the College of Biomedical Engineering and Instrument Sciences, Zhejiang University (No. 2025–7). Participants were informed about the study and could decline participation by returning a signed refusal form. Participants received monetary compensation for their participation.

**Speech and Song Recording (adults):** We collected speech and song recordings from 41 Mandarin-speaking adults (21 females; mean age = 24.0 ± 2.2 years; Table S6) in a sound-attenuated booth. Each participant spoke about 3 minutes on a self-selected topic and was allowed to prepare a written script in advance. They sang freely for about 3 minutes without instrumental accompaniment and were allowed to prepare lyrics in advance. A visual countdown timer was displayed throughout both tasks. The experimental procedures were approved by the Research Ethics Committee of the College of Biomedical Engineering and Instrument Sciences, Zhejiang University (No. 2025–6). Participants gave their informed consent and received monetary compensation for their participation.

### Sensorimotor Synchronization Experiment

**Participants:** Forty participants participated in the stimulus-synchronized vocalization experiment (23 females; mean age = 23.6 ± 1.9 years), and another forty participated in the stimulus-synchronized tapping experiment (20 females; mean age = 23.4 ± 3.0 years). In each experiment, participants were randomly assigned to either a periodic condition (n = 20) or an aperiodic condition (n = 20). In the vocalization experiment, nine participants reported more than one year of formal musical training, while in the tapping experiment, twelve participants reported more than one year of formal musical training. All participants had normal hearing and reported no history of neurological or psychiatric disorders. The study was approved by the Research Ethics Committee of the College of Biomedical Engineering and Instrument Sciences, Zhejiang University (No. 2025-006). Participants gave their written informed consent and received monetary compensation for their participation.

**Stimulus:** Each stimulus sequence consisted of a series of noise bursts. The sequence was either periodic or aperiodic. Seven mean stimulus rates were tested, i.e., 1, 2, 3, 4, 5, 6, and 7 Hz, corresponding to mean rates of one to seven noise bursts per second. Each noise burst was generated by multiplying white noise by an exponentially decaying amplitude envelope, with the decay time constant set to one-third of the target noise burst duration. In the periodic condition, the noise burst duration was fixed within each sequence, ranging from 1 s at 1 Hz to 1/7 s at 7 Hz. In the aperiodic condition, noise burst intervals were sampled from an exponential distribution whose mean matched the inverse of the target stimulus rate. To ensure that each noise burst remained audible and to avoid excessively long noise bursts, intervals shorter than 0.05 s or longer than 1.6 s were discarded and resampled. The number of noise bursts in each sequence was adjusted such that the total sequence duration just exceeded 20 s. Two sequences were generated for each condition and stimulus rate. In the periodic condition, the two sequences were identical because the inter-burst intervals were fixed. In the aperiodic condition, the two sequences were independently generated from the same exponential interval distribution.

**Procedure:** The experiment was conducted in a sound-attenuated booth. Auditory stimuli were presented binaurally via Sennheiser HD 280 Pro headphones at a self-chosen comfortable sound level. All tasks were presented using Psychtoolbox-3 in MATLAB 2022b. The auditory stimulus and the sound produced by participants were simultaneously recorded using a two-channel stereo recording at a sampling rate of 44,100 Hz, which allowed temporal alignment between the auditory stimulus and response.

In the vocalization task, participants were instructed to synchronize their vocalization as precisely as possible with the auditory stimulus: for each noise burst, they were asked to produce one syllable. After pressing the space bar to confirm readiness, the syllable to be produced in the upcoming trial (“ku”, “ti”, or “sa”) was displayed on the screen 2 s prior to stimulus onset. Participants could rest after each trial. No performance feedback was provided.

In the tapping task, participants were instructed to tap with their right index finger in synchrony with the stimulus. The procedure was identical to that of the vocal synchronization task, except that the participants responded by finger tapping instead of vocalizing.

Participants first completed a practice session in which they synchronized either vocalizations or tapping responses with three example sequences. During practice, they adjusted the sound level to ensure that the stimulus was clearly audible. Data from the practice session were not included in the analysis.

In the main experiment, participants listened to 84 sequences divided into seven blocks, each corresponding to one mean stimulus rate (1–7 Hz). Each block presented 12 sequences. In the vocalization experiment, each block contained three sub-blocks, each corresponding to one syllable (“ku”, “ti”, and “sa”), within each of which two stimulus sequences were each presented twice in randomized order (3 syllables × 2 sequences × 2 repetitions). In the tapping experiment, each block contained two stimulus sequences, each presented six times in randomized order (2 sequences × 6 repetitions). Participants were not informed about the block structure.

### Data Analysis

**Modulation spectrum analysis:** Acoustic rhythm was characterized using the modulation spectrum. The detailed procedure for calculating the modulation spectrum has been described previously (Ding et al., 2017). In brief, the sound recordings were segmented into 6-s bins and the last bin was zero-padded to 6 s (Ding et al., 2017). For each segment, the temporal envelope of sound was extracted in narrow frequency bands using a cochlear model (Yang et al., 1992). The temporal envelope in each frequency band was converted into the frequency domain using the discrete Fourier transform (DFT), and the spectrum was averaged over frequency bands. For each recording, modulation spectra were first computed for all 6-s segments and averaged using the RMS across segments. When multiple recordings belonged to the same speaker or condition, the resulting spectra were further averaged using the RMS, such that each recording contributed equally to the final estimate. The linear modulation frequency axis was resampled onto a logarithmically spaced axis (0.5–32 Hz, 200 points) using linear interpolation. The modulation spectrum of each recording (in corpus analysis) or each participant (in the sensorimotor synchronization experiment) was normalized by dividing by its maximal value, such that the peak value was scaled to 1.

For the sensorimotor synchronization experiment, we extracted the spectral envelope of the modulation spectra at different stimulus rates. In the periodic condition, we only considered the sharp spectral peaks at the stimulus rate and the first three harmonics because higher-order harmonics were substantially weaker and less reliable across participants. Across all tested stimulus rates, if multiple spectra contained peaks at the same modulation frequency, the maximal peak amplitude was retained. We linearly interpolated the peak amplitudes to obtain the spectral envelope. In the aperiodic condition, since there were no sharp spectral peaks, the spectral envelope at each modulation frequency equal the highest value across the modulation spectra across all stimulus rates.

**Linear discriminant analysis:** Linear discriminant analysis (LDA) was used to identify a one-dimensional discriminant axis that best separated speech and song modulation spectra. The LDA model was trained using adult speech and song recordings from two cross-lingual corpora (Hilton et al., 2022; Ozaki et al., 2024). In the two corpora, 96 samples were available for both speech and song. Since the sample size was limited, to avoid overfitting, we reduced the dimension of the modulation spectrum by reducing the frequency resolution. Specifically, the modulation spectrum was averaged into six octave-spaced modulation-frequency bands between 0.5 and 32 Hz (0.5–1, 1–2, 2–4, 4–8, 8–16, 16–32 Hz).

The LDA was used to derive a fixed discriminant axis rather than to perform classification. The sign of the axis was defined such that adult speech recordings had positive scores and adult song recordings had negative scores. Modulation spectra from independent corpora and additional vocalization categories were then projected onto this fixed axis to quantify their relative positions along the speech–song dimension.

**Inter-participant phase coherence spectrum:** For the sensorimotor synchronization experiment, the inter-participant phase coherence was used to quantify phase alignment across participants at each modulation frequency. The analysis was performed based on the same narrowband temporal envelopes extracted during the modulation spectrum analysis. Specifically, for each 6-s segment, the discrete Fourier transform was computed for each cochlear frequency band, yielding complex-valued coefficients at each modulation frequency. The phase angle of each DFT coefficient was extracted, and phase coherence spectrum was computed across participants at each modulation frequency as:

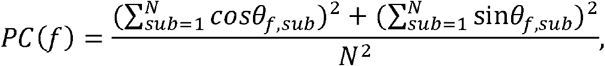

where *θ_f,sub_* denotes the phase angle at modulation frequency *f* for a given participant, and *N* denotes the number of participants.

The resulting phase coherence spectra were then averaged across cochlear frequency bands and across segments. Phase coherence spectra were computed separately for each stimulus condition, sequence, and trial repetition, and were then averaged across repetitions and sequences within the same stimulus rate, as well as across syllable conditions in the vocalization experiment. The spectral envelope of the phase coherence spectrum was estimated using the same procedure as for the modulation spectrum.

**Statistical analysis:** Statistical comparisons were performed using two-sided t tests. LDA scores were tested against zero, assessing whether a category was located on the speech or song side of the discriminant axis. Multiple comparisons were controlled using false discovery rate (FDR) correction.

## Acknowledgments

We thank Dr. Huan Luo for comments on an early version of the manuscript, Minhui Zhang, Zhenghui Sun, and Yantin Teng for data collection, and Qianxi Yu and Wenhui Sun for surveying datasets.

## Author contributions

Conceptualization, N.D.; Methodology, J.W., H.C., and N.D.; Formal Analysis, J.W. and N.D.; Resources, J.W. and N.D.; Writing – Original Draft, J.W and N.D.; Writing – Review & Editing, J.W., H.C., and N.D.; Visualization, J.W. and N.D.; Funding Acquisition, N.D.

## Supplemental information

Document S1. Figures S1–S2, Tables S1–S7.

## Data and code availability

The toddler laughter recordings and the speech and song recordings collected in this study, together with all processed data and analysis code, will be made publicly available through the Open Science Framework (OSF) upon publication (https://osf.io/ayhfq). The additional corpora analyzed in this study are publicly available, and their corresponding DOIs are provided in Table S7.

