## Supplementary Information for "Two fundamental timescales organize human vocal behavior"

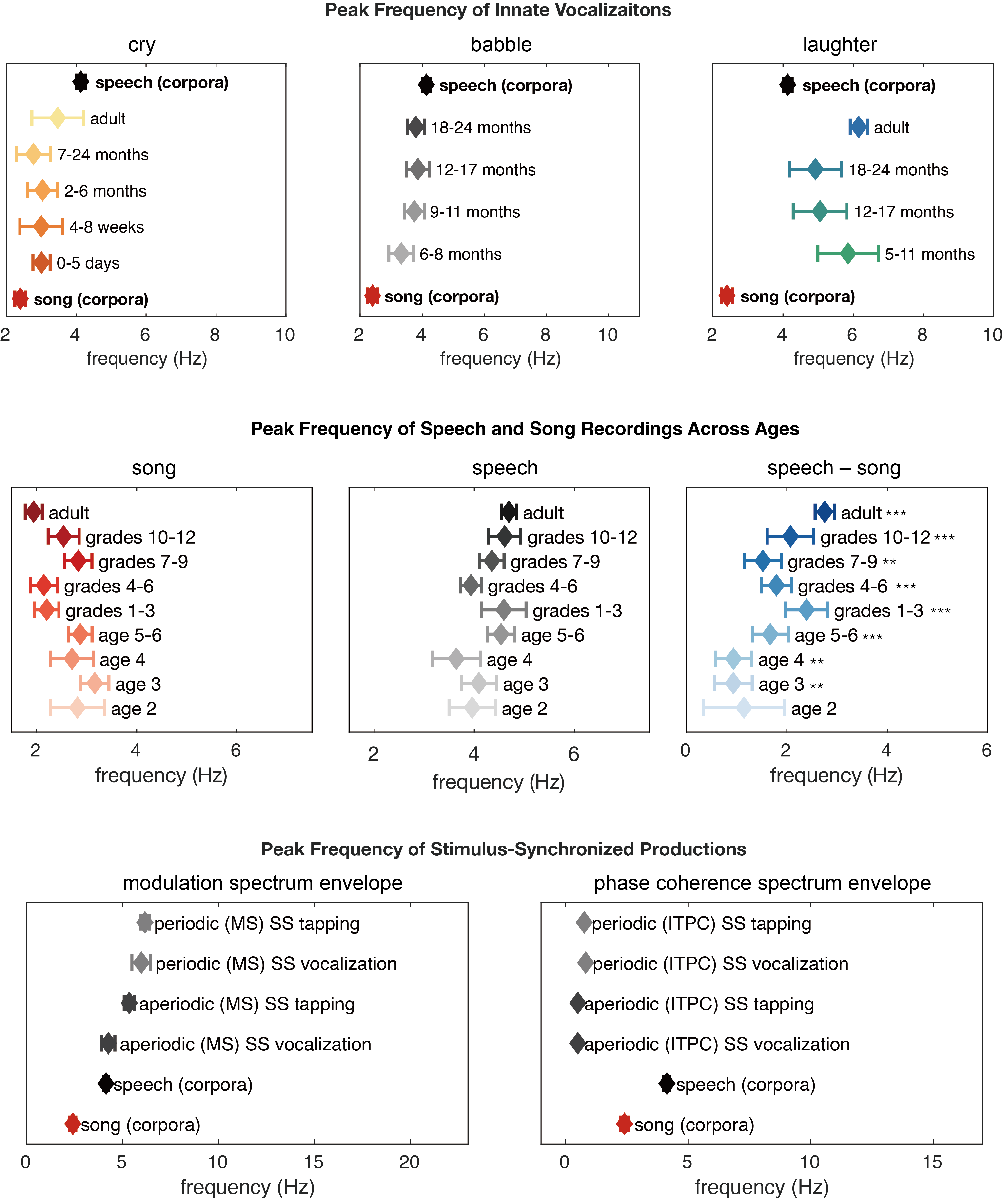
**Fig. S1.** Peak frequency of innate vocalization, speech and song recordings, and stimulus-synchronized productions. Asterisks denote significance levels of peak frequency relative to zero (*p < 0.05, **p < 0.01, ***p < 0.001; two-sided t tests with FDR correction).

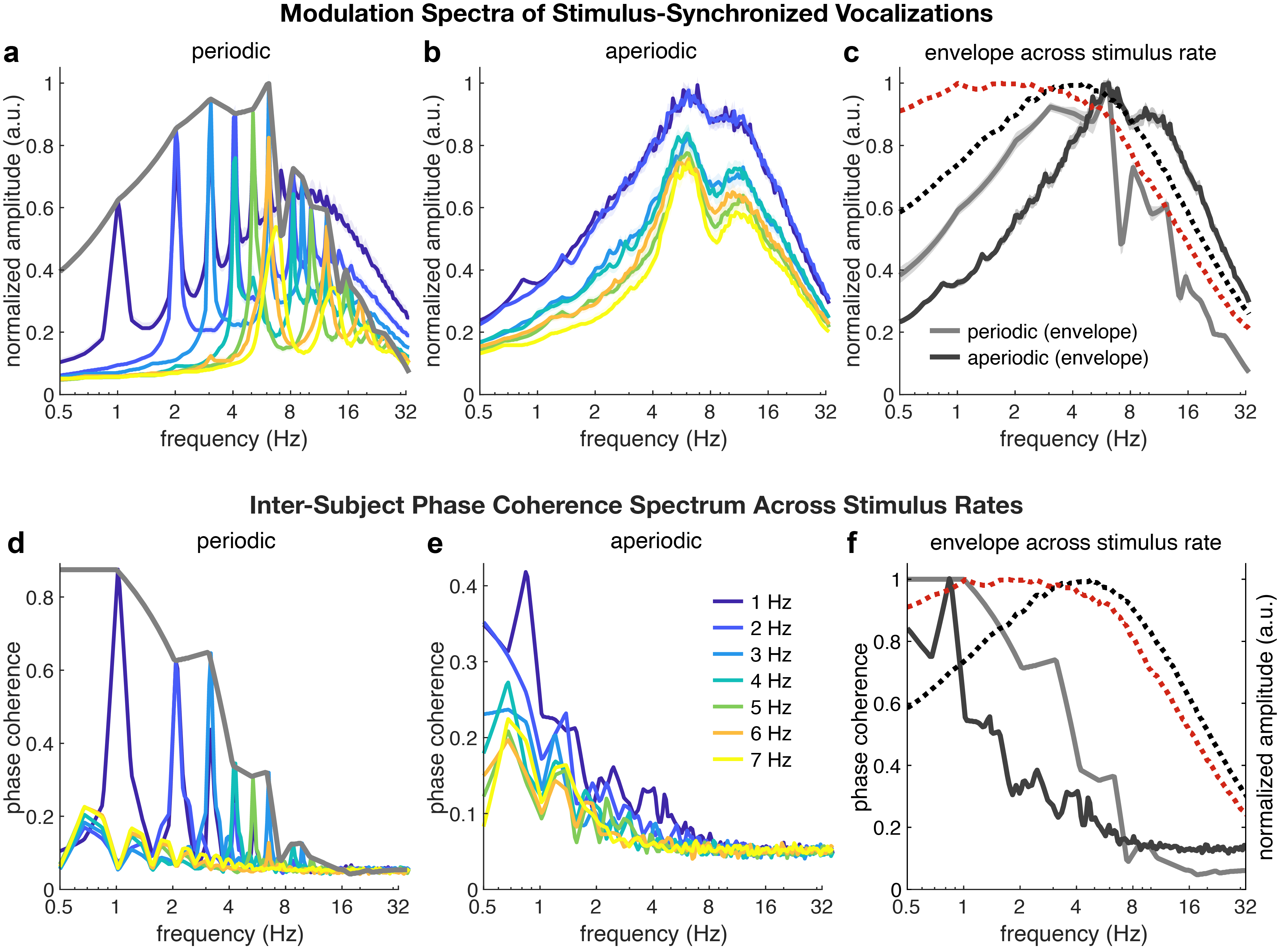

**Fig. S2.** (a-b) Modulation spectrum of stimulus-synchronized tapping at different stimulus rates (1–7 Hz) under the periodic (a) and aperiodic (b) conditions. Colors denote different stimulus rates. Shaded regions indicate ±1 SEM across participants. (c) Envelope of modulation spectra across different rates in the periodic (light gray) and aperiodic (dark gray) conditions. Shaded regions indicate ±1 SEM across participants. Reference modulation spectra of speech and song are shown as dotted lines. (d-e) Inter-participant phase coherence spectra of stimulus-synchronized tapping. (f) Envelope of phase coherence spectra across different rates.

**Table S1.** One-sample tests of LDA scores against zero across corpus

| **Corpus** | **Mean ± SE** | **t** | **df** | **P** | **P_FDR-corrected_** |
| --- | --- | --- | --- | --- | --- |
| speech (Hilton) | 2.82 ± 0.22 | 12.88 | 74 | < 0.001 | < 0.001 |
| speech (Ozaki) | 3.05 ± 0.46 | 6.64 | 20 | < 0.001 | < 0.001 |
| aishell | 3.61 ± 0.09 | 41.96 | 375 | < 0.001 | < 0.001 |
| wenet | 4.87 ± 0.15 | 33.26 | 123 | < 0.001 | < 0.001 |
| giga audiobook | 4.73 ± 0.31 | 64.44 | 263 | < 0.001 | < 0.001 |
| ted lium | 4.85 ± 0.04 | 127.46 | 645 | < 0.001 | < 0.001 |
| song (Hilton) | -2.62 ± 0.31 | -8.42 | 74 | < 0.001 | < 0.001 |
| song (Ozaki) | -3.74 ± 0.65 | -5.78 | 20 | < 0.001 | < 0.001 |
| violin | -3.09 ± 0.55 | -5.64 | 46 | < 0.001 | < 0.001 |
| viola | -4.09 ± 0.41 | -9.88 | 58 | < 0.001 | < 0.001 |
| cello | -4.89 ± 0.42 | -11.69 | 39 | < 0.001 | < 0.001 |
| bass | -5.25 ± 0.77 | -6.81 | 17 | < 0.001 | < 0.001 |
| piano | -3.71 ± 0.45 | -8.24 | 57 | < 0.001 | < 0.001 |
| guitar | -3.10 ± 0.36 | -8.63 | 44 | < 0.001 | < 0.001 |

**Table S2.** Statistical comparisons of peak frequencies across vocalization datasets

| **Comparison** | **Age group** | **Mean ± SE** | **t** | **df** | **P** | **P_FDR-corrected_** |
| --- | --- | --- | --- | --- | --- | --- |
| (A) **Innate vocalizations** | | | | | | |
| Cry  vs  Song | 0-5 d | 3.01 ± 0.05 | 2.41 | 3072 | 0.016 | 0.042 |
|  | 4-8 w | 3.00 ± 0.41 | 1.67 | 139 | 0.097 | 0.140 |
|  | 2-6 mo | 3.04 ± 0.23 | 2.27 | 194 | 0.024 | 0.053 |
|  | 7-24 mo | 2.78 ± 0.29 | 1.23 | 131 | 0.223 | 0.241 |
|  | adult | 3.47 ± 0.54 | 2.62 | 118 | 0.010 | 0.032 |
| Babble  vs  Speech | 6-8 mo | 3.33 ± 0.20 | -2.82 | 114 | 0.006 | 0.025 |
|  | 9-11 mo | 3.75 ± 0.11 | -1.57 | 120 | 0.119 | 0.155 |
|  | 12-17 mo | 3.86 ± 0.17 | -1.08 | 121 | 0.282 | 0.282 |
|  | 18-24 mo | 3.80 ± 0.08 | -1.34 | 118 | 0.184 | 0.218 |
| Laughter  vs  Speech | 5-11 mo | 5.85 ± 0.66 | 3.52 | 101 | 0.001 | 0.004 |
|  | 12-17 mo | 5.05 ± 0.56 | 2.02 | 102 | 0.046 | 0.085 |
|  | 18-24 mo | 4.92 ± 0.54 | 1.82 | 103 | 0.072 | 0.118 |
|  | adult | 6.16 ± 0.04 | 7.883 | 3480 | <0.001 | <0.001 |
| (B) **Speech and song recordings** | | | | | | |
| Speech vs song | age 2 | 1.15 ± 0.81 | 1.41 | 8 | 0.195 | 0.195 |
|  | age 3 | 0.93 ± 0.37 | 2.49 | 8 | 0.038 | 0.042 |
|  | age 4 | 0.94 ± 0.36 | 2.58 | 8 | 0.033 | 0.042 |
|  | age 5-6 | 1.67 ± 0.36 | 4.66 | 15 | < 0.001 | 0.001 |
|  | grades 1-3 | 2.39 ± 0.42 | 5.76 | 12 | < 0.001 | < 0.001 |
|  | grades 4-6 | 1.79 ± 0.30 | 6.03 | 14 | < 0.001 | < 0.001 |
|  | grades 7-9 | 1.52 ± 0.36 | 4.18 | 11 | 0.002 | 0.002 |
|  | grades 10-12 | 2.07 ± 0.46 | 4.46 | 14 | < 0.001 | 0.001 |
|  | adult | 2.75 ± 0.19 | 14.34 | 40 | < 0.001 | < 0.001 |

**Table S3.** One-sample tests of LDA scores against zero across all datasets.

| **Source** | **Age group** | **Mean ± SE** | **t** | **df** | **P** | **P_FDR-corrected_** |
| --- | --- | --- | --- | --- | --- | --- |
| (A) **Innate vocalizations** | | | | | | |
| Cry | 0-5 d | -2.46 ± 0.07 | -34.88 | 2977 | < 0.001 | < 0.001 |
|  | 4-8 w | -0.65 ± 0.48 | -1.37 | 44 | 0.179 | 0.194 |
|  | 2-6 mo | -1.06 ± 0.35 | -3.06 | 99 | 0.003 | 0.004 |
|  | 7-24 mo | -1.63 ± 0.72 | -2.28 | 36 | 0.028 | 0.034 |
|  | adult | -0.48 ± 0.99 | -0.49 | 23 | 0.632 | 0.632 |
| Babble | 6-8 mo | 1.29 ± 0.44 | 2.95 | 19 | 0.008 | 0.011 |
|  | 9-11 mo | 2.47 ± 0.23 | 10.56 | 25 | < 0.001 | < 0.001 |
|  | 12-17 mo | 2.47 ± 0.24 | 10.29 | 26 | < 0.001 | < 0.001 |
|  | 18-24 mo | 2.85 ± 0.26 | 11.04 | 23 | < 0.001 | < 0.001 |
| Laughter | 5-11 mo | 3.47 ± 0.72 | 4.81 | 6 | 0.003 | 0.004 |
|  | 12-17 mo | 3.57 ± 0.60 | 5.94 | 7 | < 0.001 | < 0.001 |
|  | 18-24 mo | 3.78 ± 0.59 | 6.39 | 8 | < 0.001 | < 0.001 |
|  | adult | 4.75 ± 0.04 | 126.85 | 3373 | <0.001 | <0.001 |
| (B) **Speech and song recordings (children and adult)** | | | | | | |
| Speech | age 2 | 2.49 ± 0.81 | 3.06 | 8 | 0.016 | 0.016 |
|  | age 3 | 3.29 ± 0.66 | 5.00 | 8 | 0.001 | 0.001 |
|  | age 4 | 3.37 ± 0.59 | 5.71 | 8 | 0.001 | 0.001 |
|  | age 5-6 | 4.29 ± 0.39 | 11.04 | 15 | < 0.001 | < 0.001 |
|  | grades 1-3 | 3.41 ± 0.41 | 8.25 | 12 | < 0.001 | < 0.001 |
|  | grades 4-6 | 3.65 ± 0.38 | 9.57 | 14 | < 0.001 | < 0.001 |
|  | grades 7-9 | 4.62 ± 0.35 | 13.26 | 11 | < 0.001 | < 0.001 |
|  | grades 10-12 | 3.52 ± 0.28 | 12.80 | 14 | < 0.001 | < 0.001 |
|  | adult | 4.30 ± 0.21 | 20.24 | 40 | < 0.001 | < 0.001 |
| Song | age 2 | 0.14 ± 0.75 | 0.19 | 8 | 0.851 | 0.914 |
|  | age 3 | -0.74 ± 0.62 | -1.19 | 8 | 0.267 | 0.400 |
|  | age 4 | -0.66 ± 0.92 | -0.71 | 8 | 0.496 | 0.638 |
|  | age 5-6 | -0.06 ± 0.52 | -0.11 | 15 | 0.914 | 0.914 |
|  | grades 1-3 | -1.71 ± 0.61 | -2.83 | 12 | 0.015 | 0.034 |
|  | grades 4-6 | -3.21 ± 0.53 | -6.07 | 14 | < 0.001 | 0.001 |
|  | grades 7-9 | -1.50 ± 0.66 | -2.28 | 11 | 0.044 | 0.078 |
|  | grades 10-12 | -2.23 ± 0.56 | -3.95 | 14 | < 0.001 | 0.004 |
|  | adult | -4.30 ± 0.32 | -12.49 | 40 | < 0.001 | < 0.001 |
| Speech minus  Song | age 2 | 2.34 ± 1.17 | 2.01 | 8 | 0.079 | 0.079 |
|  | age 3 | 4.03 ± 0.48 | 8.40 | 8 | < 0.001 | < 0.001 |
|  | age 4 | 4.29 ± 0.94 | 4.28 | 8 | 0.003 | 0.003 |
|  | age 5-6 | 4.35 ± 0.65 | 6.66 | 15 | < 0.001 | < 0.001 |
|  | grades 1-3 | 5.12 ± 0.65 | 7.87 | 12 | < 0.001 | < 0.001 |
|  | grades 4-6 | 6.85 ± 0.53 | 12.87 | 14 | < 0.001 | < 0.001 |
|  | grades 7-9 | 6.12 ± 0.76 | 8.01 | 11 | < 0.001 | < 0.001 |
|  | grades 10-12 | 5.75 ± 0.61 | 9.41 | 14 | < 0.001 | < 0.001 |
|  | adult | 8.33 ± 0.34 | 24.84 | 40 | < 0.001 | < 0.001 |
| (C) **Stimulus-synchronized envelopes of modulation spectrum** | | | | | | |
| vocalization | periodic | 2.68 ± 0.58 | 4.60 | 19 | < 0.001 | < 0.001 |
|  | aperiodic | 5.84 ± 0.44 | 13.38 | 19 | < 0.001 | < 0.001 |
| tapping | periodic | 4.67 ± 0.40 | 11.61 | 19 | < 0.001 | < 0.001 |
|  | aperiodic | 8.46 ± 0.23 | 36.75 | 19 | < 0.001 | < 0.001 |

**Table 4.** Summary for vocalization corpora

| **Vocalization Type** | **Corpus / Source** | **Language** | **N speaker** | **Speaker Duration**  **(s, M ± SD)** | **Total Duration (min)** |
| --- | --- | --- | --- | --- | --- |
| adult cry | Deeply | Korean | 24 | 5.4 ± 0.5 | 2.1 |
| adult laughter | Vocalsound | Multilingual | 3,505 | 4.2 ± 1.9 | 247.2 |
| adult laughter | Deeply | Korean | 25 | 5.2 ± 0.5 | 2.1 |
| babble | Phonbank | Multilingual | 27 | 13747.1 ± 10033.8 | 6186.2 |
| infant cry | Donate-a-cry | Multilingual | 195 | 8.6 ± 7.7 | 27.8 |
| infant cry | EnesBabyCries | Multilingual | 75 | 1503.5 ± 1169.1 | 1879.3 |
| toddler laughter | this study | Mandarin | 27 | 119.3 ± 107.4 | 53.7 |
| song | Song-Speech | Multilingual | 75 | 55.5 ± 33.2 | 69.4 |
| song | Human-vocalization | Multilingual | 21 | 444.0 ± 324.2 | 155.4 |
| speech | Song-Speech | Multilingual | 75 | 48.2 ± 33.1 | 60.3 |
| speech | Human-vocalization | Multilingual | 21 | 706.2 ± 624.7 | 247.2 |

**Table S5.** Toddler participant demographics

| **Age group** | **N** | **age (month)** | **Females** | **Duration (s)** |
| --- | --- | --- | --- | --- |
| 5-11 months | 8 | 9.00 ± 1.93 | 5 | 57.12 ± 27.14 |
| 12-17 months | 11 | 13.82 ± 1.47 | 5 | 71.36 ± 81.05 |
| 18-24 months | 9 | 21.11 ± 2.32 | 5 | 45.44 ± 28.89 |

**Table S6.** Summary for speech and song recording (children and adult)

| **Age Group** | **N Speaker (female)** | **Age (years, M ± SD)** | **Speech Speaker Duration**  **(s, M ± SD)** | **Speech Total Duration (min)** | **Song Speaker Duration**  **(s, M ± SD)** | **Song Total Duration (min)** |
| --- | --- | --- | --- | --- | --- | --- |
| age 2 | 11 (5) | 2.6 ± 0.2 | 98.3 ± 65.9 | 18.0 | 77.6 ± 32.8 | 14.2 |
| age 3 | 9 (4) | 3.6 ± 0.2 | 141.3 ± 104.8 | 21.2 | 93.1 ± 52.6 | 14.0 |
| age 4 | 9 (5) | 4 ± 0.0 | 98.3 ± 55.7 | 14.8 | 105.8 ± 54.7 | 15.9 |
| ages 5-6 | 16 (9) | 5.5 ± 0.5 | 182.5 ± 140.7 | 48.7 | 91.9 ± 40.8 | 24.5 |
| grades 1-3 | 13 (8) | 8.3 ± 0.9 | 125.9 ± 66.4 | 27.3 | 126.9 ± 111.7 | 27.5 |
| grades 4-6 | 15 (8) | 10.3 ± 0.8 | 132.5 ± 59.0 | 33.1 | 153.0 ± 59.0 | 38.3 |
| grades 7-9 | 12 (9) | 13.6 ± 1.1 | 109.1 ± 36.4 | 21.8 | 110.2 ± 45.4 | 22.0 |
| grades 10-12 | 15 (13) | 16.9 ± 0.4 | 105.3 ± 24.5 | 26.3 | 113.3 ± 23.6 | 28.3 |
| adult | 41 (21) | 24.0 ± 2.2 | 193.5 ± 17.0 | 132.2 | 176.6 ± 23.7 | 120.6 |

**Table S7.** Key Resources Table

| **REAGENT or RESOURCE** | **SOURCE** | **IDENTIFIER** |
| --- | --- | --- |
| Software and algorithms | | |
| Python 3.13.2 | Python Core Team | https://python.org/ |
| Matlab R2022b | MathWorks | https://ww2.mathworks.com/ |
| Other | | |
| Processed data and code | This paper | https://osf.io/srfe5/ |
| Deeply | OpenSLR | https://www.openslr.org/99/ |
| Vocalsound | Gong et al., 2022 | https://github.com/YuanGongND/vocalsound |
| Phonbank | Talkbank | https://talkbank.org/phon/ |
| Donate-a-cry | Gabor Veres | https://github.com/gveres/donateacry-corpus |
| EnesBabyCries | Lockhart-Bouron et al., 2023 | https://osf.io/ru7na/ |
| Song-Speech | Ozaki et al., 2024 | https://osf.io/mzxc8/overview |
| Human-vocalization | Hilton et al., 2022 | https://zenodo.org/records/5525161 |
